# Asynchrony rescues statistically-optimal group decisions from information cascades through emergent leaders

**DOI:** 10.1101/2022.04.05.487127

**Authors:** Andreagiovanni Reina, Thomas Bose, Vaibhav Srivastava, James A. R. Marshall

## Abstract

It is usually assumed that information cascades are most likely to occur when an early but incorrect opinion spreads through the group. Here we analyse models of confidence-sharing in groups and reveal the opposite result: simple but plausible models of naïve Bayesian decision-making exhibit information cascades when group decisions are synchronous; however, when group decisions are asynchronous, the early decisions reached by Bayesian decision makers tend to be correct, and dominate the group consensus dynamics. Thus early decisions actually rescue the group from making errors, rather than contribute to it. We explore the likely realism of our assumed decision-making rule with reference to the evolution of mechanisms for aggregating social information, and known psychological and neuroscientific mechanisms.

## 1 Introduction

Information cascades, where individuals follow others’ decisions regardless of self-sourced evidence, are usually assumed to occur in asynchronous decision-making, in which early decisions tend to be incorrect, and dominate the decision dynamics so that the group decision is incorrect. Previous work assumed cascades to happen only when the first responding individual exerts disproportionate influence on other group members [1, 2, 3, 4]. The converse assumption would be that synchronous group decision-making mechanisms offer the best protection from information cascades.

Here we explore the optimal pooling of information in synchronous and asynchronous group decision-making mechanisms. A standard assumption in behavioural ecology, psychology, and neuroscience is that individuals apply optimal probabilistic computational rules where possible (*e.g*. [5, 6, 7, 8, 9, 10]). If optimal computation is infeasible, it is argued that rules that approximate optimal computations in typically-encountered scenarios will be used. Similarly, in behavioural ecology and psychology, research focuses on the optimal pooling of information by groups (*e.g*. [11, 12]).

In evolutionary terms, the neurological mechanisms to process asocial environmental information must have developed earlier than sociality appeared. Thus, we assume that when group living began, evolution led to the adaptation of pre-existing Bayesian heuristics [7] to also process social information. We study the implications of such assumptions on group decisions in two scenarios: collective detection of an instantaneous signal with synchronous interaction among individuals, and continuous environmental sampling with asynchronous interaction. Our analysis shows that for the synchronous case, in which there are no early decisions, decision-making is unstable and negative information cascades are observed. In the asynchronous case, however, early decisions tend to be correct and lead to positive information cascades. This observation is the opposite to the usual assumption that early decisions are erroneous and lead to negative information cascades, showing how group leaders can spontaneously emerge for the benefit of collective decisions.

### 1.1 Problem formulation

We study the problem of a single-shot collective decision in which *N* individuals pool information to make a decision on the correct state of the world *S*. We assume the choice is binary, that is, there are two possible states of the world *S* ∈ {*S*^+^, *S*^−^}. We assume that each state of the world has a prior probability, *P* (*S*^+^) and *P* (*S*^−^). We assume that the cost matrix for classifications is symmetric, *i.e*. the cost of an error, as well as the reward for the correct classification, is the same for either state of the world. We consider two types of collective decisions: *signal detection* and *sequential sampling* (Fig. 1).

**Figure 1:**
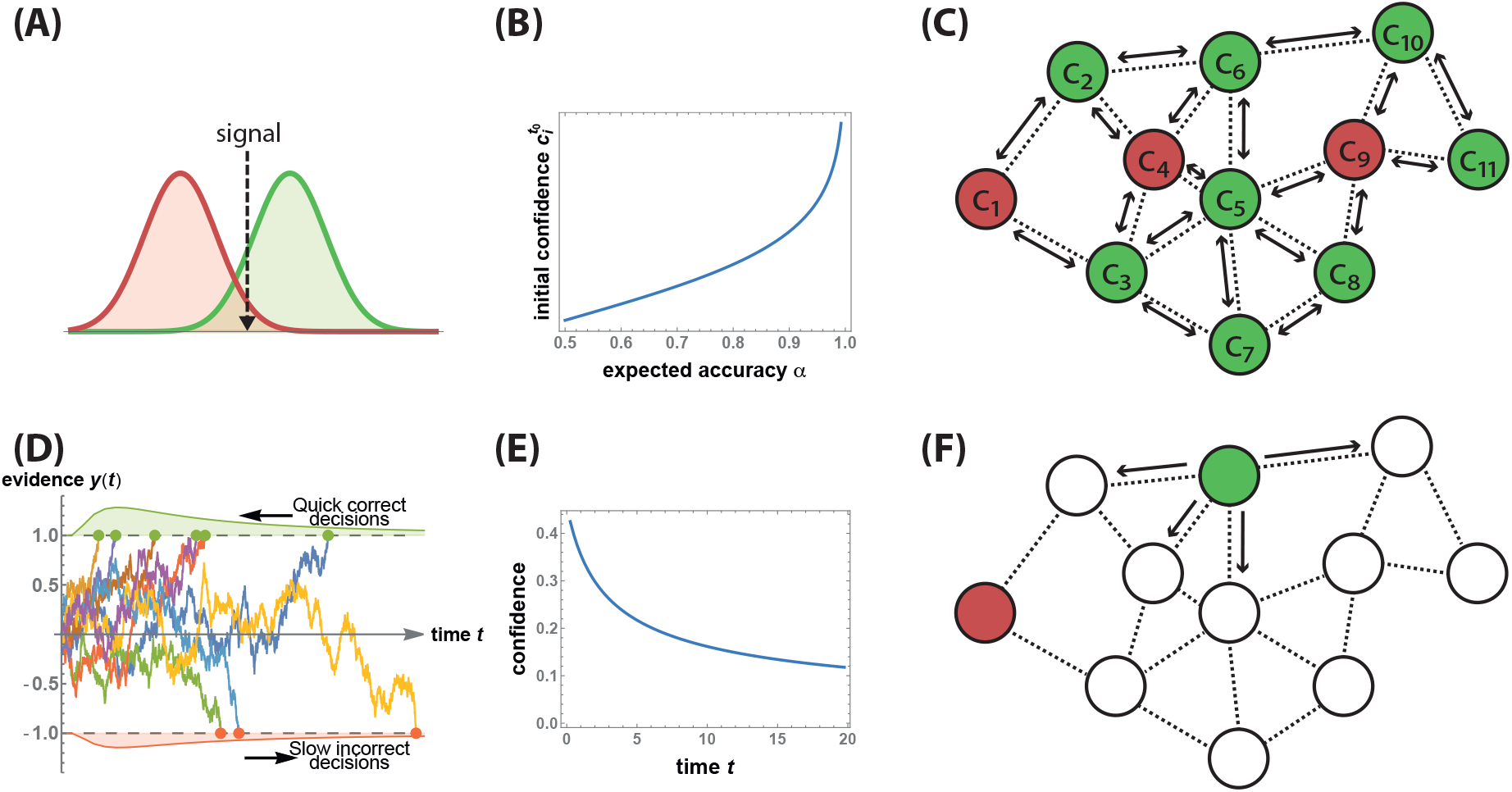
We consider two types of collective decisions—(A-C) signal detection and (DF) sequential sampling—characterised respectively by synchronous and asynchronous social interactions. **(A)** An instantaneous event at time *t*_0_ produces a signal that all individuals estimate and compare with a threshold to make a decision between the red and green alternatives (signal detection theory). **(B)** We assume that each agent has an estimate of its accuracy (*e.g*. through previous experience), with which it can estimate its confidence 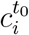 as the log-odds ratio, Eq. (1), Ref. [11]. **(C)** Individuals synchronously exchange options and confidence *c*_*i*_ with their nearest neighbours (*i.e*. information spreads on a random geometric graph [13]), and, in order to reach a consensus decision, they update their opinions by locally-optimal Bayesian integration of confidence-weighted votes (Weighted Bayes Consensus rule). The arrows indicate bidirectional synchronous interactions, the colours are the individuals’ opinions. **(D)** In sequential sampling, each individual optimally integrates noisy evidence from the environment until it has enough information to make a decision. This process is modelled as a Drift Diffusion Model (DDM). The graphics shows examples of DDM trajectories for drifts sampled from a random distribution biased towards the correct decision as positive drift. The expected decision time is shorter for correct decisions (positive threshold) and longer for incorrect decisions (negative threshold), because, as indicated in [14], errors are in most cases caused by low drift-diffusion ratios which take longer, on average, to reach the decision threshold than DDMs with high drift-diffusion ratios which lead in most cases to correct decisions. **(E)** When the individual does not know its DDM’s drift but can only estimate its expected sampling ability, its confidence (computed with Eq. (4)) is high when the accumulated evidence hits the decision threshold early (a quick decision is a proxy of higher DDM’s drift-diffusion ratio, in agreement with neurological mechanisms [15]) and low when it hits the threshold late. **(F)** An individual (node) only communicates once it makes a decision, which it communicates to its neighbours (in the graphics, the green node with one-way communication arrows; the red node has reached its decision earlier and does not continue communicating).

In signal detection, at time *t*_0_, individuals are exposed to a signal emitted by an instantaneous event, which they categorise as *S*^+^ or *S*^−^ (Fig. 1A). Through optimal signal detection theory, individuals compare the estimated signal with a threshold as described in Sec. 1.2.1. Therefore, each individual *i*, at time *t*_0_, has an independent opinion 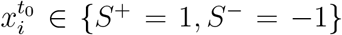 on the true state of the world *S* and the relative confidence 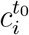 on the accuracy of its opinion (Fig. 1B). Every individual *i* repeatedly exchanges its opinion and confidence with its neighbours *M*_*i*_ defined by a communication network topology 𝒢, which can be static or time-varying. Individuals, at each synchronous social interaction, update their opinion and confidence in order to determine the correct state *S* (Fig. 1C).

In sequential sampling, each agent integrates evidence from the environment over time in order to correctly classify the state of the world. As in neuroscientific studies [16], the statistically optimal process of evidence integration is represented as a *Drift Diffusion Model* (DDM), [17, 18], which describes the evolution over time of the indi-vidual *i*’s decision evidence *y*_*i*_(*t*) as a biased Brownian motion process that is governed by two terms: the drift *A*_*i*_ and the diffusion *W* (Fig. 1D). The former term models the evidence integration towards the correct decision, while the latter term models the noise in the integration process (implemented as a Wiener process with standard deviation *σ* equal for every agent). Individuals integrate evidence *y*_*i*_(*t*) until one of the two thresholds *z*^+^ *>* 0 or *z*^−^ *<* 0 has been reached (*i.e*. we model a free-response scenario). The individual’s decision corresponds to the sign of the crossed threshold, or equivalently of the integrated evidence, *i.e*. 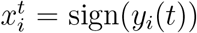). Individuals making a decision communicate it to their neighbours *M*_*i*_ on 𝒢 (Fig. 1E-F), who combine the received information with their accumulated evidence as described in Sec. 1.2.2.

### 1.2 Weighted Bayes Consensus

We formulate how naïve Bayes-optimal individuals can employ the statistically optimal Bayes’ rule [19] to update their opinion 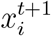 and confidence 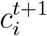 from their neighbours’ opinions in the two scenarios considered, collective signal detection and collective sequential sampling. We describe updates as naïve-Bayes because they neglect correlations in social information [20].

#### 1.2.1 Collective signal detection

In signal detection, individuals form an opinion in favour of either *S*^+^ = 1 or *S*^−^ = −1 by comparing the estimated signal with a threshold specific to each agent (Fig. 1A). We assume that each agent *i* has an estimate of its accuracy *α*_*i*_ in determining the true state of the world. This information can have been acquired by the individual, for example through previous experience and decisions in the same environment. In this experimental scenario, at time *t*_0_, each individual synchronously makes an independent estimate 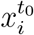 of the world’s state. Following optimal signal detection theory [11], each agent *i* can also compute its confidence 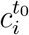 as the log-odds ratio

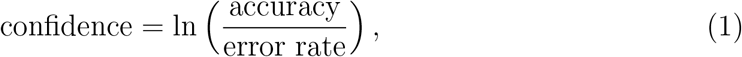

where the error rate is the complementary probability of being correct, *i.e*. 1−accuracy, thus 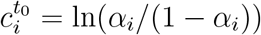, Fig. 1B.

After the individual decisions, every iteration *t > t*_0_ the individuals share with each other their opinion 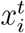 and confidence 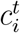, and use the received information to update their new opinion 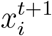 and new confidence 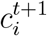 (Fig. 1C). Statistically optimal individuals compute the new aggregate opinion 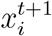 following optimal confidence weighting theory presented in [21, 22, 11] as

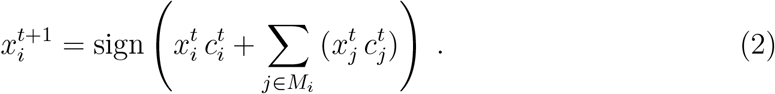

This leaves, however, the problem of how individuals update their confidence in their new opinion. We start by noting that the neighbours’ confidences can be used to derive the neighbours’ accuracies (or, equivalently, vice versa). Assuming that all agents update their confidence through the same computation, the inverse of the confidence computation of Eq. (1) gives the accuracy of each neighbour (see *Materials and Methods*). We label this update rule as Weighted Bayes Consensus, and in *Materials and Methods* we show it can be reduced to linear summation

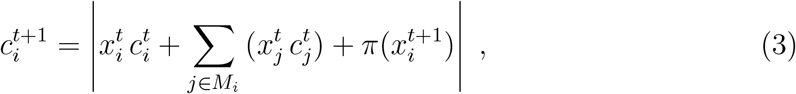

where the operator | ‐|is the absolute value, and 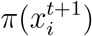 is the log of the prior ratio in favour of 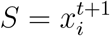, *i.e*. 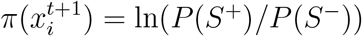 for 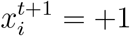, and the reciprocal of the log argument for 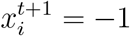. Therefore, the Weighted Bayes Consensus rule is a simple linear update rule for both opinion (Eq. (2)) and confidence (Eq. (3)).

#### 1.2.2 Collective sequential sampling

In the sequential sampling scenario, individual *i* makes a decision 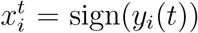 when the integrated evidence *y*_*i*_(*t*) reaches the threshold *z*^+^ *>* 0 or *z*^−^ *<* 0 in favour of the positive or negative world state hypotheses, respectively, at time *t* (Fig. 1D). The thresholds are optimally set as a function of the priors *P* (*S*^+^) and *P* (*S*^−^), and the cost matrix, following [16] (for details see Text ST1 in the supplementary material). Note that the thresholds are set to an equal and fixed value for all individuals, as every individual has the same knowledge at the beginning of the decision making process. We assume that agents know the integration noise *σ*, the cost matrix, and the world state priors *P* (*S*^+^) and *P* (*S*^−^), which are the same for the entire population, in agreement with previous theory [7]. Individuals do not know their drift *A*_*i*_—which represents the individual’s accuracy in sampling the state of the world [16]—but they know the random distribution from which the drifts’ magnitudes, *Ã*_*i*_, are sampled (assuming no systematically misinformed individuals, the sign of *A*_*i*_ is always equal to the correct state *S*). In other terms, individuals know the group accuracy distribution, but do not know the accuracy of any specific individual.

An individual *j* communicates its decision 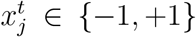 to its neighbours *M*_*j*_ once—when its integrated evidence *y*_*j*_(*t*) reaches either threshold (*z*^+^, *z*^−^), see Fig. 1F. The information about another individual reaching threshold is additional evidence that the neighbours can use during their continuous evidence integration. Therefore, an individual *i* that receives the neighbour’s decision 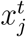 at time *t* and has not yet made a decision (*i.e. z*^−^ *< y*_*i*_(*t*) *< z*^+^), integrates 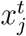 as a ‘kick’ *k* into its evidence accumulator *y*_*i*_(*t*). The optimal size of this kick corresponds to the neighbour’s confidence in its decision which depends both on the quantity of integrated evidence and the integration time (Fig. 1E). In general, quick decisions are considered as an indication of high confidence (due to a high drift-diffusion ratio *Ã*_*i*_*/σ*), conversely slow decisions are likely to be influenced by high levels of noise (low *Ã*_*i*_*/σ*); see Fig. 1D and Ref. [14]. Note that there is no difference if the decision-maker computes its own confidence (*k*) and sends this information, or if every agent infers *k* once receives a neighbour’s decision. Assuming identical thresholds in the population and simultaneous start of evidence integration, an agent receiving a neighbour’s decision has information on both the integration time *t* (*i.e*. communication time) and on the integrated quantity 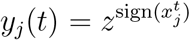 (*i.e. z*^+^ for 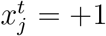 and *z*^−^ for 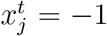). Therefore the optimal kick size, for example assuming 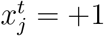, is

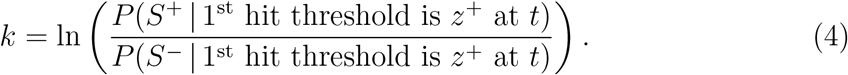

Again, applying Bayesian theory [19] we obtain

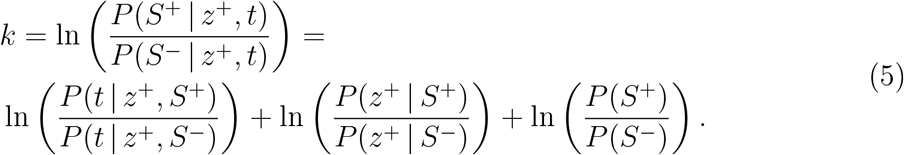

The three terms of the rhs of Eq. (5) respectively are the log-odds of the first passage time of the DDM through *z*^+^ at *t*, the log-odds of hitting *z*^+^ before *z*^−^, and the log-odds of the prior on the states of the world. The precise DDM parameters are unknown to the individual thus, as proposed in [15], the individual averages the probability for any possible DDM weighted by the prior probability of such DDM parameters to manifest. See *Materials and Methods* for the detailed derivation and Fig. 1E for a graphical illustration of Eq. (5).

More sophisticated agents could aggregate social information with more advanced computation that uses absence of decision from neighbours as informative data [23]. Similarly, an individual could refine the computation of Eq. (5) by observing the social network on its neighbours and treating differently the case in which the neighbour makes a decision based solely on its personal information or after receiving social information [20]. Such nuanced calculations are likely to be unrealistic to be implemented in the brain, hence in our study, we assume naïve individuals that neglect previous social interactions. Signals from neighbours making their decisions are treated independently, thus each neighbour is implicitly considered as the frist decider, consistent with the naïve-Bayes assumption. We base our assumptions on the argument that mechanisms for optimal evidence integration of asocial cues have been co-opted to the social case, without any refinement.

### 2 Results

We quantified the effect of the proposed rules on a group of *N* individuals that cooperate through social signalling with each other. In both tested scenarios—collective signal detection and sequential sampling—we assumed individuals communicate on a partially connected network, that is, each individual *i* has a limited number of neighbours *M*_*i*_ *< N*. We conducted our tests on Random Geometric Graphs (RGG) [13], which are constructed by locating the *N* nodes at uniform random locations in a unit square, and connecting two nodes when their Euclidean distance is smaller than *δ*. The value of *δ* determines the average degree connectivity *κ*—that is, the average number of neighbours each individual has. We chose to study interaction on a RGG topology as it closely relates to systems embedded in a physical environment, thus matching the characteristics of several biological systems. Results for other types of network topologies are reported in the supplementary material.

#### Synchronous updates lead to negative information cascades

As noted above, the naïve Bayes-optimal signal detection rule, Weighted Bayes Consensus, gives linear updating of both decisions (Eq. (2)) and confidence (Eq. (3)). In *Materials and Methods* we show that such linear updating of confidence leads to an unstable process on the agent network; this means that decisions will be precipitated more rapidly than in stable processes, but at the expense of accuracy. In Fig. 2 we numerically compare the speed and accuracy of Weighted Bayes Consensus against the Belief Consensus algorithm [24], through which every individual, by iteratively averaging weighted opinions over its neighbourhood, computes the weighted mean of the entire population (find a detailed description of the algorithm in Sec. 5.3). On the one hand, Weighted Bayes Consensus is the locally optimal solution, as individuals apply the Bayes-optimal signal detection rule on information locally available at each moment; on the other hand, Bayes Consensus is the globally optimal solution, as after a number of iterations every individual computes the global weighted average (Eq. (2) computed on every member), which corresponds to the optimal solution to the collective signal detection problem [11]. In both cases, optimality is defined in terms of accuracy only, assuming naïve individuals. In the *Discussion* we consider the relevance of these algorithms for natural systems; for now we note that, as group heterogeneity varies, a speed-accuracy trade-off is described (Fig. 2). Compared with the Belief Consensus algorithm, the Weighted Bayes Consensus is dominated on group accuracy but takes on average a shorter time to reach consensus. This comparison makes it possible to appreciate the effect of the unstable dynamics of the Weighted Bayes Consensus in contrast to the slower but stable dynamics of the Belief Consensus algorithm. Fig. 2 also shows that the group improves in collective accuracy with increasing heterogeneity *σ*_*α*_, as a consequence of higher mean individual accuracy (see also Fig. SF1 in the supplementary material).

**Figure 2:**
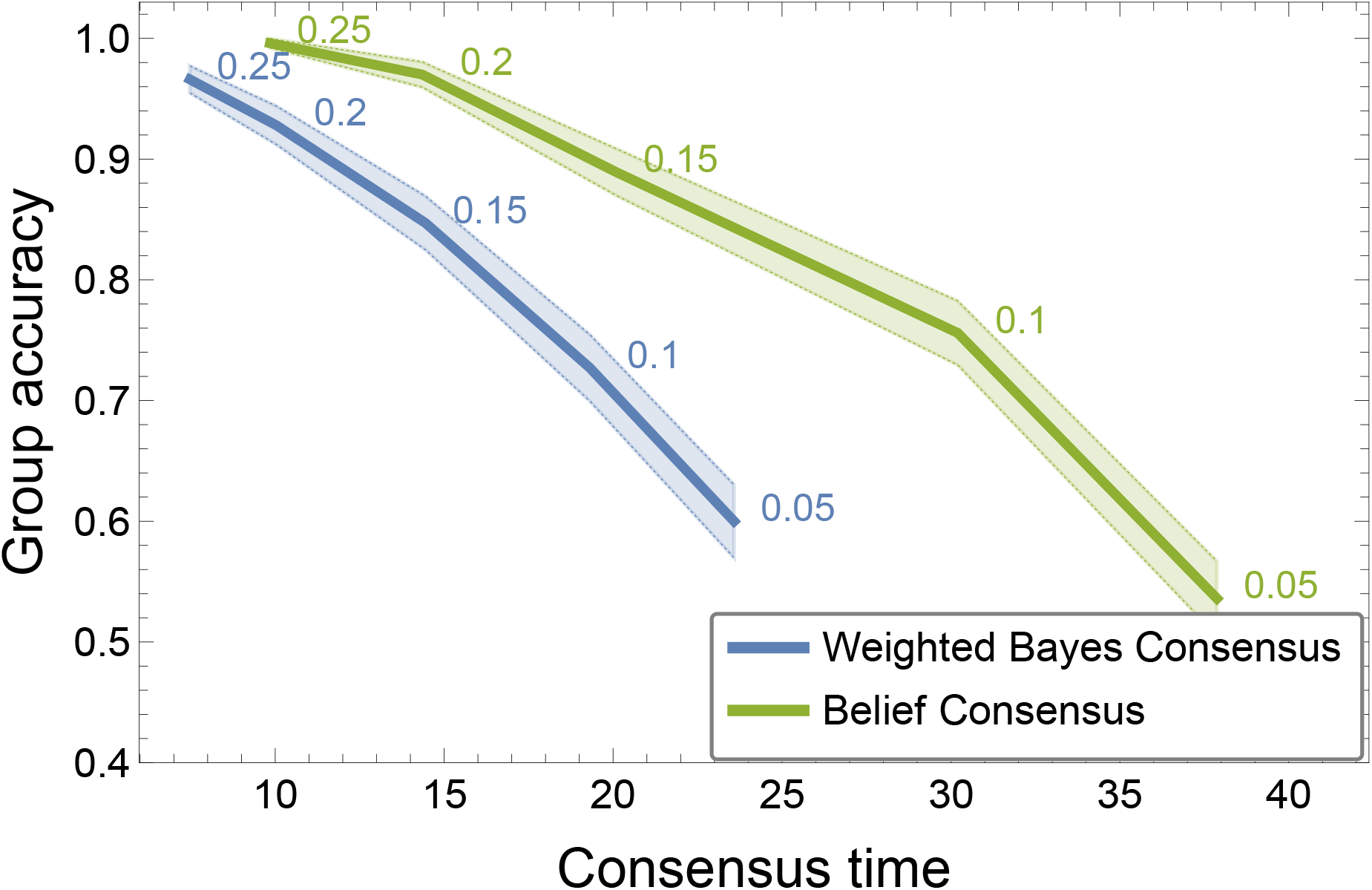
Via synchronous updates of individuals’ confidence through the Weighted Bayes Consensus rule—which neglects correlation of social information—the group reaches a consensus in less time than the optimal strategy (‘Belief Consensus’). However, quick runaways can lead to erroneous decisions, as shown by a lower group accuracy. Here we show the results for 10^3^ simulations of *N* = 50 individuals that have individual accuracy *α* drawn from a normal distribution 𝒩 (*μ*_*α*_ = 0.5, *σ*_*α*_) (flipping to 1 − *α* when *α <* 0.5), and varying heterogeneity *σ*_*α*_ (shaded areas are 95% confidence intervals). The Weighted Bayes Consensus rule (WBC, blue lines) has a lower group accuracy than the Belief Consensus algorithm (BC, green lines) however is quicker (heterogeneity level *σ*_*α*_ is indicated next to the curve; group accuracy is computed as the proportion of runs with unanimous agreement for *S*^+^).

#### Asynchronous updates prevent negative information cascades through the emergence of informed leaders

In collective sequential sampling, individuals can be assumed to incur a cost that is a linear function of *ω*_*e*_ for erroneous decisions (assuming that correct decisions incur no cost) and *ω*_*t*_ for the time taken to make their decision. This can be defined according to the Bayes Risk [25, 16], enabling agents to set optimally their decision thresholds in order to minimise expected cost [26, 16] (see Text ST1 in the supplementary material). For collective decisions in sequential collective decision-making we find that, contrary to the synchronous case, larger numbers of information cascades are triggered by the best decision-makers (Fig. 3). This is because, on average, the best individuals are expected to reach their decision threshold quicker than others (Fig. 1(d), Ref. [14]). Such early signals cause a larger response than delayed decisions (Fig. 3(a)). The resulting effect is that the best individuals—those that are more accurate because they have a higher signal to noise ratio *Ã/σ*—more often trigger a cascade of decisions in the group (Fig. 3(b)), and the best decision-makers’ cascades are typically larger than the ones triggered by the inferior individuals (Figs. 3(c) and SF6). Therefore, we observe that on average the best decision-makers have the highest influence on the group, acting as emergent group leaders as a direct consequence of a combination of psychological and neuroscientific mechanisms [16, 14, 15].

**Figure 3:**
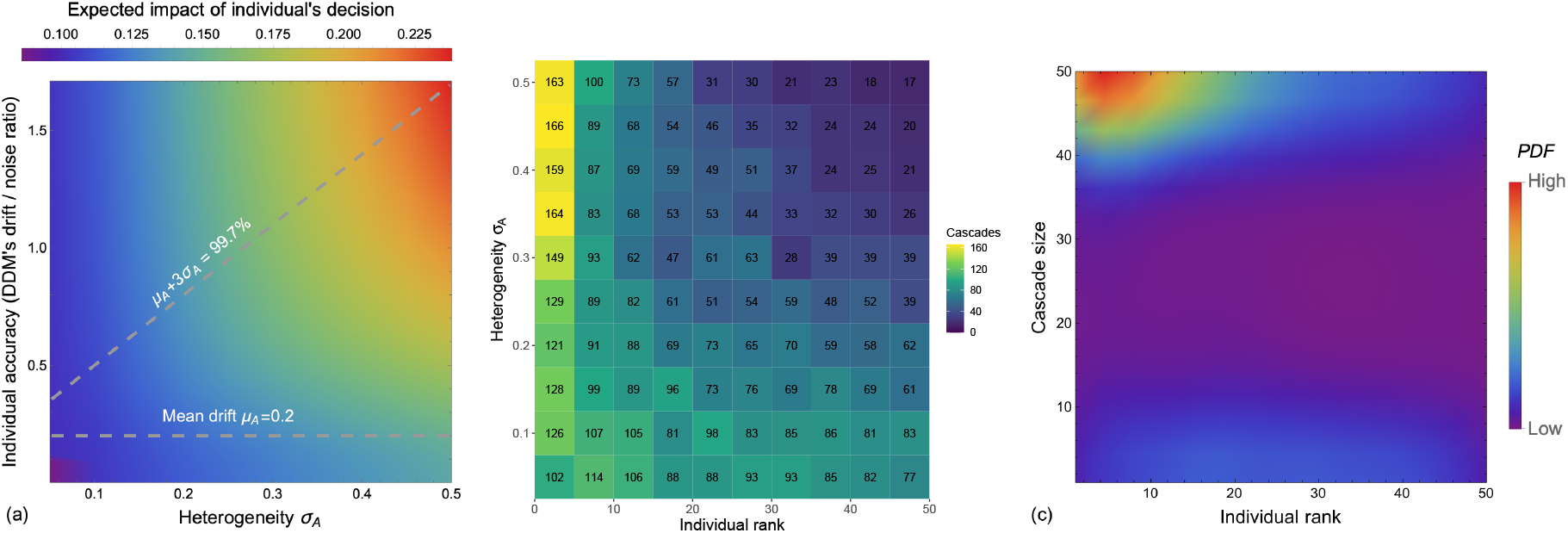
Emergent leaders from psychological and neuroscientific mechanisms. (a) We report the expected impact of an individual decision on its neighbours. In diverse groups, the most accurate individuals are expected to have a large impact on others. We computed the expected decision time for each DDM with noise *σ* = 1 and drift sampled from the normal distribution 𝒩 (*μ*_*A*_, *σ*_*A*_) (with *μ*_*A*_ = 0.2, and *σ*_*A*_ varied on the x-axis, with 3*σ*_*A*_ indicated with dashed lines). The threshold *z* is set to optimise the Bayes Risk with costs *ω*_*t*_ = 1 and *ω*_*e*_ = 100. Higher drifts are expected to reach the threshold earlier [16], and earlier reactions are considered a sign of higher confidence [15]. For each case, we visualise the kick size *k* (Eq. (5)) at the expected decision time, normalised by the threshold *z, i.e*. the colour indicates *k/z*. Therefore larger values bring the individual closer to its decision threshold. (b) In diverse groups, the best individuals— that is, with higher drift/noise ratio, *Ã/σ*—more often trigger a cascade. We sort (on the x-axis) the individuals in decreasing order of drift/noise ratio and report the number of cascades each individual triggers. We count as a cascade the triggering of a sequence of at least *N/*10 decisions. The results are from 500 simulation runs of a group of *N* = 50 individuals communicating on a sparse network (connected random geometric graph with average degree *κ* = 10), and with drift sampled from 𝒩 (*μ*_*A*_ = 0.2, *σ*_*A*_). In more homogenous groups (low *σ*_*A*_), cascades are almost equally likely to be triggered by any individual. Instead, in highly heterogenous groups (high *σ*_*A*_), cascades are predominantly caused by the best individuals. (c) The most accurate individuals trigger the largest cascades. We show the probability density function for each individual, sorted in decreasing drift/noise ratio order, to trigger a cascade of different sizes (on the y-axis). The PDF is computed from 500 runs for the case of *σ*_*A*_ = 0.5 and the same parameters of panel (b). Thus, in summary, leaders emerge in heterogeneous groups as their decisions are followed (a) strongly, (b) more frequently, and (c) by a larger portion of the population.

### 2.1 Model comparison

The goal of this study is to show that, contrary to common intuition [1, 2, 3, 4], early decisions can have a beneficial impact on the collective dynamics by triggering positive information cascades, even in populations of naïve-Bayesian agents, whereas in absence of temporal ordering among decisions (synchronous scenario), naïve-Bayesian agents can frequently suffer negative information cascades. Despite being biologically unrealistic (as further discussed in Sec. 4), the collective dynamics in the synchronous scenario can be rescued by a simple change in the individual behaviour, by averaging neighbours’ opinions rather than summing them (Belief Consensus algorithm). Our analysis also explains the causes of our results. In particular, we compute the mathematical stability and instability of the synchronous scenario systems when there is perpetual integration of social information and we indicate how confidence can be inferred from the decision speed based on known neuroscientific mechanisms [15, 16].

Here, we explicit similarities and differences in the two models and in the assumptions on which the two scenarios are based. Both scenarios describe how individuals integrate social information in oder to improve their own world’s estimate. Both scenarios are also based on the same assumptions that individuals are naïve because they neglect correlations in social information and locally integrate social evidence through Bayes-optimal rules according to the information they have access to. Hence, the considered strategies are optimal in terms of accuracy on the presumption of naïve individuals; we further discuss the biological relevance of our assumptions in Sec. 4. Notwithstanding the strong similarities, the two scenarios differ in terms of the environmental information the individuals integrate, how and when they communicate with one another, and, consequently, in the rules to combine their opinion and confidence with the ones of their neighbours (see Fig. 1). As a consequence of such differences, the performance of the two scenarios cannot be compared directly, rather we show how (mis-)information cascades have a different impact on the two scenarios. Comparing quantitatively the speed-accuracy results of both scenarios is impractical. In fact, in Fig. 2, we only analyse the runs of the signal detection scenario that reached unanimous agreement; however, a condition of unanimous consensus is rare in the sequential sampling scenario, because individuals do not change their decision once they reached a threshold. Although there is no consensus, Figs. SF2(a-b) show that, in sequential sampling, a large majority of the group makes correct decisions, more frequently than in an asocial condition. The objective of our analysis is to show the negative impact of quick runaways in the synchronous signal detection scenario (Fig. 2), and explain that the situation is the opposite in the asynchronous sequential sampling scenario where the large majority of information cascades are triggered by individuals making correct early decisions (Fig. 3). These results generalise to different network topologies and all tested parameters, as shown in Figs. SF3, SF4, SF5, and SF6.

## 3 Previous work

Collective decision making in groups of individuals that update their opinion beliefs has been widely investigated, commencing with the seminal model of DeGroot [27]. Collective decision-making models have been investigated in the social sciences in the form of social learning and in engineering as consensus-averaging algorithms. We briefly review previous relevant approaches.

### Non-Bayesian social learning

A large amount of work has investigated social learning [1, 2, 28, 3, 29, 30, 31, 32, 4, 33, 34, 35, 36, 37, 38, 39, 40, 41] in which individuals update their beliefs with a Bayes-optimal rule that assumes correlation neglect, also referred to as non-Bayesian social learning. The correlation neglect assumption is that individual agents do not take account of the fact that incorporating neighbours’ social information with their own iteratively leads to correlated information. Instead when individuals know the full network topology they can apply the actual Bayes-optimal update rule, as in [42, 43, 44], or approximations of it [45, 46], although even with full information doing so may be computationally prohibitive. In studies of non-Bayesian social learning, various aspects have been analysed, such as the conditions for polarisation of the population [35, 41], or how information cascades can be the result of non-Bayesian update of local beliefs [1, 2, 3, 4]. Studies showed how correlation neglect can improve the performance of voting systems [47] or lead to the formation of extremists [37, 48, 41, 38]. In these studies, individuals sequentially make their rational decision based either on all previous individual decisions [1, 2, 3] or only the previous [4]. As more individuals make the same decision, the probability the next individuals will ignore their personal opinion and follow the social information becomes higher [49]. Individuals neglect correlation of information and the ordering of previous decisions, which can have determining effects on the collective dynamics as shown in [50]. In our work, we do not externally impose the ordering of votes, rather we test both synchronous simultaneous voting and asynchronous signalling with the ordering determined by the environmental sampling dynamics.

### Consensus averaging algorithms

As a form of social learning, consensus averaging algorithms allow the nodes of a network, each having a numeric value, to compute in a decentralised way the average of all these values. Therefore, through these decentralised algorithms, each agent on a sparse graph can converge on the same average confidence value. The Belief Consensus algorithm [24] uses a linear function, while other averaging algorithms employ nonlinear [51, 52, 53, 54] or heterogenous functions [55]. The advantage of consensus averaging algorithms is a guarantee of convergence in a relatively small number of timesteps. Consensus-averaging opinion dynamics models have also shown unbounded increases in individual agent confidence, leading to the formation of extremists in populations [56, 57, 58, 59, 60, 61, 62].

### Optimal evidence accumulation

The dynamics of a network of optimal evidence accumulators has been investigated in the form of coupled Drift Diffusion Models (DDMs) [63] in which each accumulator can access the state of its neighbours prior to reaching its own decision. Accessing the internal state of other agents is biologically implausible and accordingly, in our work, neighbours only share their decision when the decision threshold is reached. A similar recent study, [20], has derived the theory to allow optimal decision makers, modelled as DDMs, to update their evidence based on neighbours’ decisions (once the neighbour’s evidence reaches the decision threshold). However, this work makes the biologically-unrealistic assumptions that agents are truly Bayes-optimal, and do not use correlation neglect as a computational shortcut. These assumptions require the agents to know the complete communication topology 𝒢 in order to compute ‘second-order’ evidence integration over the behaviour of the neighbours of neighbours. The calculations rapidly become very intricate. Additionally, in integrating only neighbours’ decisions, but not the time take to reach those decisions, the agents modelled by [20] neglect an important information source, which we incorporate into our model. In agreement with previous analysis [64], our model predicts that the mean collective cost (computed from decision time and errors) decreases by increasing group heterogeneity and group connectivity (see Fig. SF2 in the supplementary material).

## 4 Discussion

We have shown analytically that for synchronous decisions locally-optimal Bayesian integration of weighted votes, in order to reach a group decision, is described by an unstable linear dynamical system in which erroneous decisions dominate. As shown numerically in comparison to an existing linear consensus algorithm with guaranteed convergence, this results in faster decisions but at the expense of group decision accuracy. In contrast, when decisions are asynchronous early decisions tend to be correct and hence, through confidence-signalling, leaders can spontaneously emerge from the best informed members of a group, and precipitate fast and accurate group decisions. That animal groups exploit the skills of the best individuals has already been observed [65, 66], however in our analysis, group leaders emerges from social interactions as the consequence of applying confidence mechanisms from neuroscience [16, 15] to social dynamics.

Our results can be interpreted through the lens of ‘information cascades’ in decisionmaking groups of humans and other animals, in which early erroneous information is assumed to dominate (*e.g*. [67, 68, 69, 70]). In contrast to this accepted view, however, negative information cascades occur when decisions are synchronous, so there are no ‘early’ decisions, but the move to asynchronous decisions actually results in early decisions being correct more often than incorrect and, correspondingly, leads to positive rather than negative information cascades on average. Our predictions are consistent with the empirical observations of collective decision making in fish [71], in which the first fish making a decision is generally no less accurate than later fish. Despite standard theory on sequential choices suggests the first decision maker should perform worse, empirical results [71] and our analysis indicate the opposite: early responses are the consequence of having access to better information, and thus acting on that information sooner. Correct and early responders can be individuals with better abilities to discriminate between environmental stimulus and noise, either due to systematic higher capabilities [65, 66] or to occasional access to a better information source (*e.g*. due to a better position) [71]. While our model is based on confidence mechanisms from neuroscience [16, 15], we do not exclude the possibility that in some species decision order may also be determined by individual traits, such as boldness or impulsivity [72].

Our analysis assumes that optimal rules for asocial information integration may have been co-opted to social scenarios where they are non-optimal, since they neglect correlated information. In the literature, correlation neglect has been studied under different names, such as “bounded rationality” [33], “imperfect recall” [73, 40], “persuasion bias” [30, 35], or “naive inference” [38]. Such correlation neglect has been observed in experiments with humans, which are cognitively advanced organisms that could in principle solve the correlation problem but still neglect to do so [74, 75, 76, 77]. Thus, since natural selection acts at the level of the individual rather than the group [78] our results may help provide a normative explanation for such apparently non-adaptive behavioural outcomes. Indeed, evidence of maladaptive social information leading to suboptimal group decision-making has been reported in several species via empirical observations [79, 69, 80, 81, 82, 70, 83], and theoretical models [84, 85].

As noted, a superior solution to decision-making under correlation neglect exists for the synchronous decision case in the form of the Belief Consensus algorithm which averages rather than sums information from neighbours. Changing to use this method of evidence integration would be straightforward even for selection acting on individuals within groups, since the behavioural selection is at the level of the individual, and membership of a group in which decisions are reached more effectively is individually advantageous. If evolutionary stable, this change of strategy would globally improve collective decision-making, but would not contradict our results as interactions are synchronous and there are no early decisions. It is important noting that, regardless of which strategy has a higher selective advantage, in any case, the synchronous decision model is a very unrealistic abstraction of biological reality. In contrast, for the more realistic scenario of asynchronous decisions, avoiding correlation neglect is informationally and computationally very demanding [74, 75, 76, 77], hence the heuristic of applying naïve-Bayesian evidence integration to social information is highly plausible, and under this reasonable assumption early decisions tend to precipitate positive rather than negative information cascades, in contradiction to previous assumptions.

## 5 Materials and Methods

Our method applies Bayes’ rule [19] to specify how the individual *i* should compute a Bayes-optimal integration of its *M*_*i*_ neighbours’ opinions to update its opinion 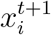 and its confidence 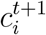.

### 5.1 Integrating neighbours’ confidence into collective signal detection

Each individual *i* communicates to its neighbours *M*_*i*_ its opinion 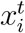 and its confidence 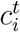. Assuming all individuals use the same computation of Eq. (1) to derive their confidence, its inverse gives the accuracy of each neighbour:

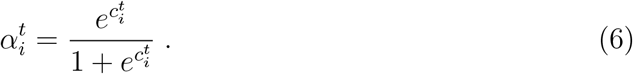

Given the set of received votes 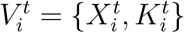 as the combination of received opinions 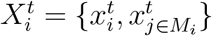 at time *t* and the set of accuracies 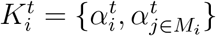 from Eq. (6), the agent *i* can compute its confidence from the probability that the aggregated opinion 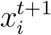 is correct (*i.e*. the true state of the world *S* ∈ *{S*^+^, *S*^−^*}* is equal to the individual’s opinion 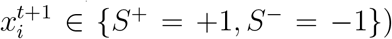. The new confidence 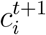 corresponds to the log-odds of being correct rather than incorrect:

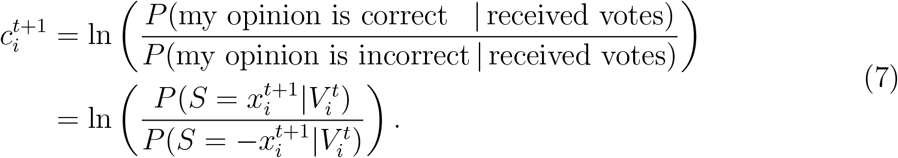

Neglecting information correlations, a statistically optimal individual can compute the probability 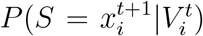 that the aggregated opinion 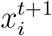 is correct given the re-ceived votes 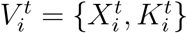 using Bayes’ rule as

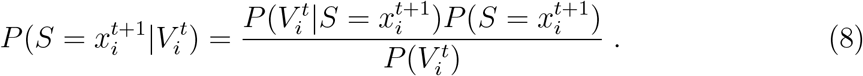

where the probability of observing the votes 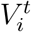 assuming 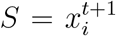 corresponds to a simple multiplication of probabilities as

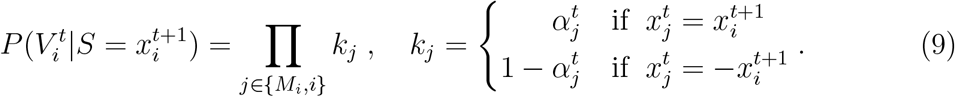

From Eqs. (6) and (9) we have that if 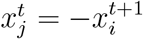 then

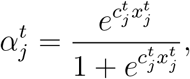

and if 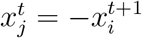, then

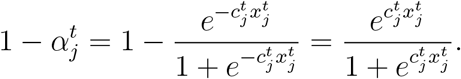

Therefore for Eq. (9), irrespective of the sign of 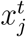 we have that

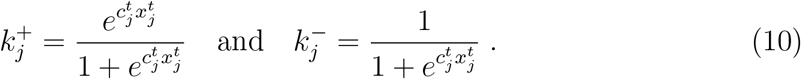

Using the above simplification, the update of Eq. (7) becomes

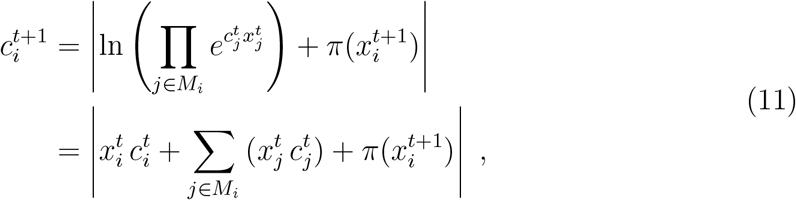

where Eq. (11) corresponds to Eq. (3) in the main text.

### 5.2 Sequential sampling scenario

In the sequential sampling scenario, an individual *i* that is integrating evidence and receives at time *t* a decision 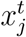 from its neighbour *j*, updates its evidence variable *y*_*i*_(*t*) by *k* which it computes with Eq. (5). The first two terms of this equation are the log-odds of the first passage time of the DDM through the threshold for 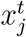 at *t* and the log-odds of hitting the threshold for 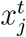 before the one for 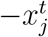. If, without loss of generality we assume 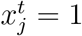, the first-passage time through *z*^+^ is computed, following the results of [86], as

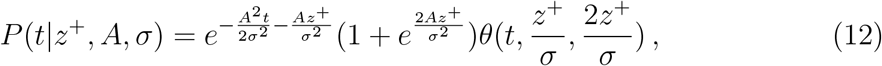

where the function *θ*(*t, u, v*) is defined as

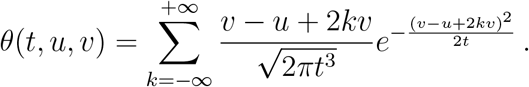

Instead, the probability of hitting *z*^+^ before *z*^−^ is

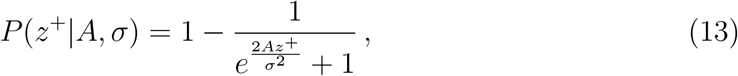

as from [16].

The individual does not know the drift rate but only knows the random distribution from which the drift is sampled. Therefore the individual integrates all possible drifts over the given random distribution and Eq. (4) can be rewritten as

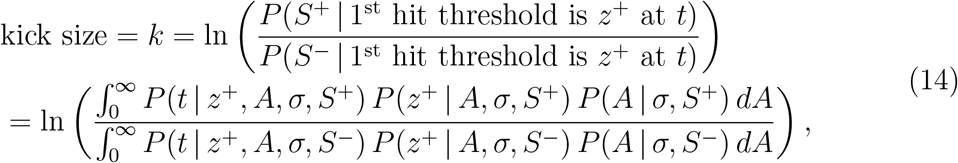

Recall that *S*^+^ determines the sign of *A*, and therefore

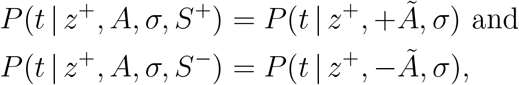

and equivalently applies for Eq. (13).

### 5.3 Analytical comparison

We compare the dynamics of the proposed Weighted Bayes Consensus rule and the linear consensus averaging algorithm from the literature, Belief Consensus [24]. Belief Consensus is a decentralised algorithm which allows each agent on a sparse graph to converge on the same average value [24]. Each agent *i* runs the algorithm by repeatedly integrating information received from its neighbours *M*_*i*_. The algorithm implements linear updates that provably converge on global consensus in a finite number of timesteps. The algorithm is defined as

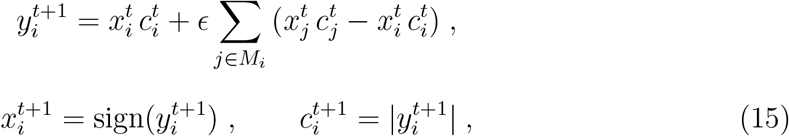

where 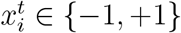 is the option selected by agent *i* at time *t*, 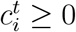 is its confidence at that time defined according to Eq. (1), *M*_*i*_ are its neighbours, and *E* is a parameter. Given the Laplacian matrix *L* of the connectivity graph 𝒢, in order to guarantee convergence the parameter *ϵ* must be chosen so that (*I* − *ϵL*) is a doubly stochastic matrix (where *I* is the identity matrix of appropriate dimensions). Metropolis-Hastings matrices are among the state-of-the-art techniques to compute *E* in a decentralised fashion using the local neighbourhood only [87].

We focus on the dynamics of 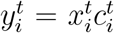, where 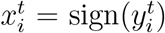 and 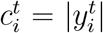, as from

Eqs. (15) and (11). Let ***y***^*t*^ be the vector of 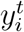’s. Given a graph 𝒢 without self-loops, we denote its adjacency matrix by *A*. Using this notation, we can rewrite Eq. (11) as

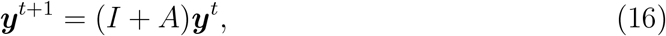

where *I* is the identity matrix of appropriate dimensions. Similarly, we can rewrite the Belief Consensus as

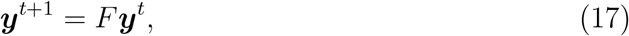

where *F* is a row stochastic matrix.

Both the Belief Consensus (Eq. (15)) and the Weighted Bayes Consensus (Eq. (16)) are linear dynamical systems. It is known that if the underlying graph is connected, the dynamics of Eq. (17) converge to the average of the initial values of ***y***, *i.e*., to 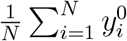, where *N* is the number of agents [24]. This convergence is a consequence of the fact that for a connected graph, the matrix *R* has one eigenvalue at 1 with associated eigenvector **1**_*N*_, and all remaining eigenvalues are inside a unit disk centered at the origin. In the context of hypothesis testing, the aggregate log-odds (log-odds of all agents pooled together) is compared against a single threshold. In this sense, the dynamics of Eq. (17) yields the correct statistic at each node which can be compared against the correct threshold, which in our case is zero, (*i.e*. we need simply determine the sign of 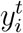). Note that for the consensus ***y***^*t*^ is always bounded.

The dynamics of Eq. (16) replace the action of averaging with the neighbours with the action of simply adding the value of the neighbours to the current agent’s value. Note that the dynamics of Eq. (16) are unstable for most graphs, *i.e*., the value of ***y***^*t*^ grows unboundedly. The agents ignore this instability as the opinion 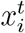 is determined only by the sign of 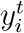. The underlying idea is that the projection of the initial condition onto the eigenvector associated with the largest eigenvalue will dominate after a small initial transient, and will be indicative of the sign of the average pooled statistic. However, the eigenvector associated with the largest eigenvalue of *I* + *A* is not the ones vector **1**_*N*_ except for regular graphs. Except for regular graphs, the dominant mode of ***y***^*t*^ will not be associated with the average statistic and will not yield the desired accuracy, but since ***y***^*t*^ will grow exponentially, it will be very quick in reaching a region in which the sign of 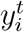 will be stable.

Note that because we are only interested in the sign of the average of the initial conditions, we could also leverage instability to reach quicker decisions in the case of the dynamics of Eq. (17). In Eq. (15), we could destabilise Eq. (17) by introducing the tuneable parameter *ϵ >* 0 as follows:

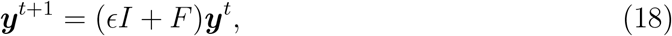

where *I* is the identity matrix of appropriate dimensions. This dynamics will have the dominant eigenvalue of 1 + *ϵ*, and associated eigenvector **1**_*N*_. Hence the dominant (unstable) mode will correspond to the average of initial conditions.

## Acknowledgments

We thank Naomi Leonard for helpful discussions. This study was partially funded by the European Research Council (ERC) under the European Union’s Horizon 2020 research and innovation programme (grant agreement number 647704). AR also acknowledges support from the Belgian F.R.S.-FNRS, of which he is a Chargé de Recherches.

## Data availability

Data and relevant code for this research work are stored in GitHub: https://github.com/DiODeProject/DecisionsOnNetworks and have been archived within the Zenodo repository: https://doi.org/10.5281/zenodo.7032373.

## Authors’ contribution

JARM proposed the models. JARM, AR, and VS mathematically analysed the models. AR numerically analysed the models. All authors wrote the manuscript.

## Conflict of interest statement

The authors declare no conflict of interest.

## Supplementary material

### ST1. Optimal DDM’s thresholds

Given the cost matrix and the priors *P* (*S*^+^) and *P* (*S*^−^), it is possible to compute the optimal thresholds that minimise the Bayes Risk cost function [16]. We assume a symmetric cost matrix where both positive and negative (either true or false) predictions have the same cost. In our simulation, a correct decision (true positive and true negative) has zero cost, while an incorrect decision (false positive and false negative) has the same cost *ω*_*e*_ = 100. The cost *ω*_*t*_ = 1 indicates unitary cost for each unit of time taken to make the decision. With these assumptions, we compute the Bayes Risk (BR) in the same way as previous work [25, 26, 16], as the weighted sum of the decision time (DT) and the error rate (ER),

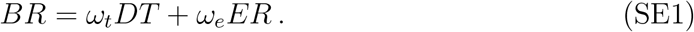

While in the paper we present our theory for the generic case, for simplicity in our simulation we set equal neutral priors *P* (*S*^+^) = *P* (*S*^−^) = 0.5. In this way, symmetric cost matrix and priors allow us to use equal thresholds *z*^+^ = − *z*^−^ = *z*. As indicated in [16] (in Eq.(46)), in this case, the Bayes Risk is minimised by the threshold *z* satisfying the following condition:

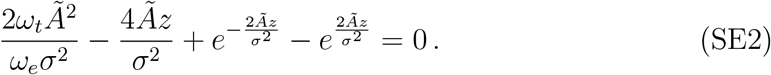

This equation does not have closed form solutions for *z*, therefore we approximate the optimal *z* numerically [16].

Because the individuals do not know the magnitude of their drift *Ã*, and instead they only know the random distribution from which *Ã* is drawn, the individuals set their threshold by integrating all possible drifts over the given random distribution, computing the threshold that minimises the BR for each drift and weighting each threshold for the probability density of that drift.

**Figure SF1:**
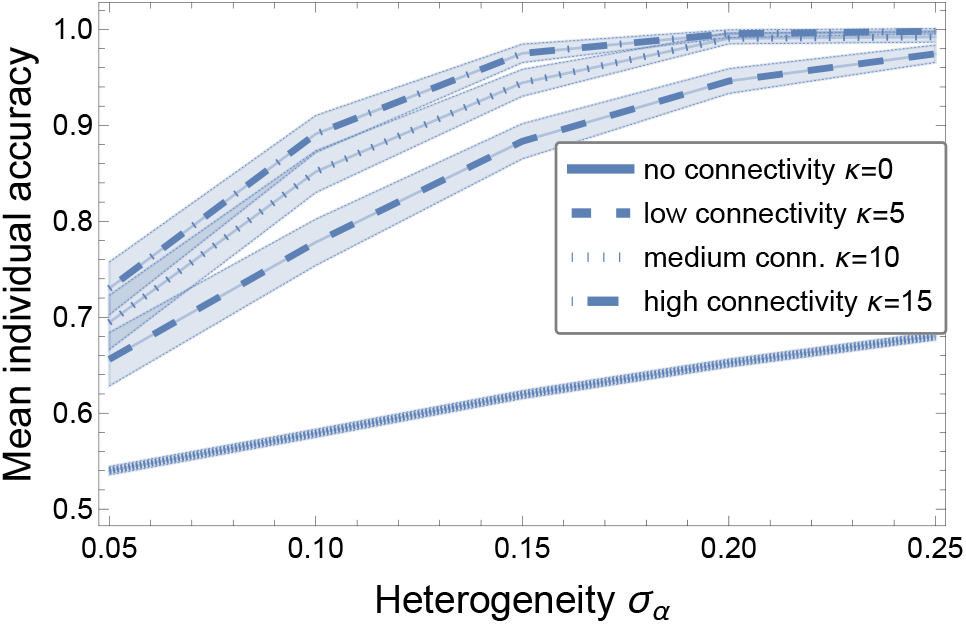
We show the results for 10^3^ simulations of *N* = 50 individuals that make synchronous updates of their confidence through Weighted Bayes Consensus rule and have individual accuracy *α* drawn from a normal distribution 𝒩 (*μ*_*α*_ = 0.5, *σ*_*α*_) (flipping to 1 − *α* when *α <* 0.5), and varying heterogeneity *σ*_*α*_ (shaded areas are 95% confidence intervals). Increasing the connectivity, we observe a significant increase of the mean individual accuracy (data for low individual accuracy *μ*_*α*_ = 0.5).

**Figure SF2:**
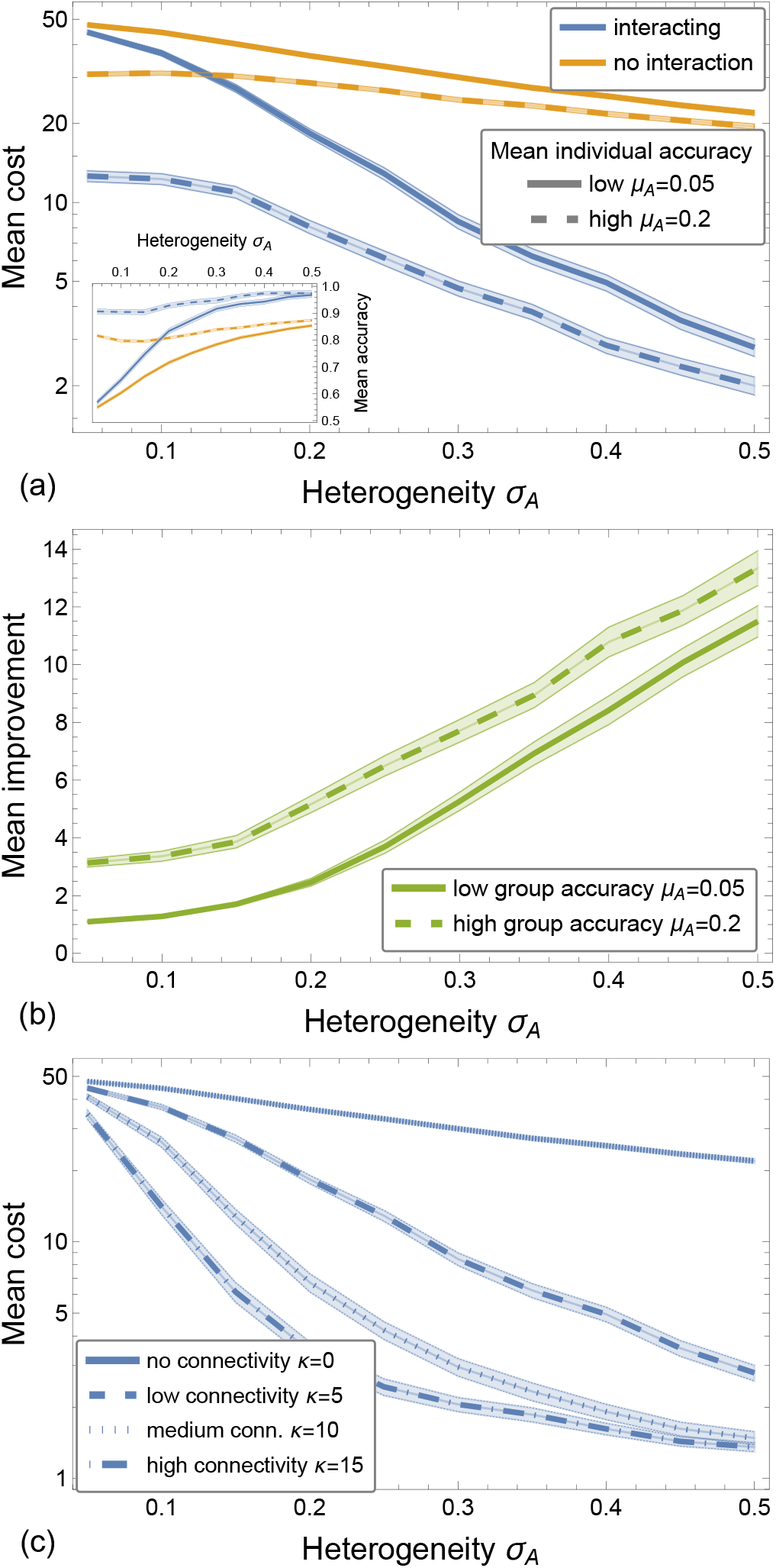
In collective sequential sampling, the benefits of social interactions are larger with increased group heterogeneity and higher connectivity. We show the mean of 500 simulations of *N* = 100 individuals that behave as a DDM, each with noise *σ* = 1 and drift sampled from 𝒩 (*μ*_*A*_, *σ*_*A*_) (considering only positive drifts), with costs *ω*_*t*_ = 1 and *ω*_*e*_ = 100, and varying heterogeneity *σ*_*A*_ (shaded areas are 95% confidence intervals). (a) Through social interactions (blue lines), the mean cost decreases more than when individuals make independent decisions (orange line). The mean cost decreases with increased heterogeneity. The inset shows the mean accuracy as the expected proportion of individuals correctly categorising the state of the world. (b) The improvement due to interaction is larger with increased group heterogeneity. The improvement is computed as the ratio between the cost incurred by the group with social interactions over the asocial group, for each of the 500 independent simulation runs. (c) Increasing the connectivity, we observe a significant decrease of the mean cost (data for low group accuracy *μ*_*A*_ = 0.05).

**Figure SF3:**
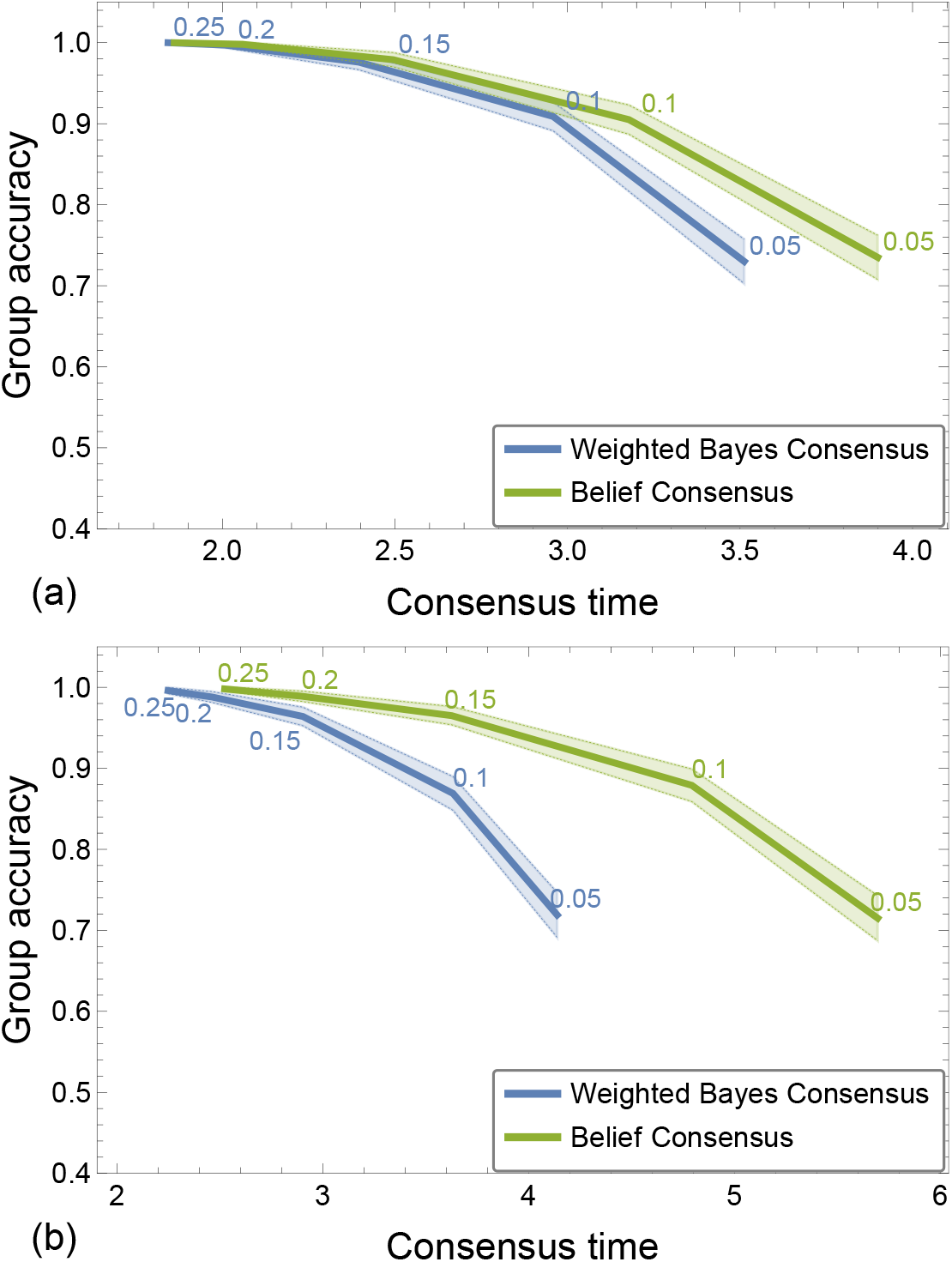
The results of Fig. 2 in the main text generalise to other types of network topologies. We report the results for the same parameters reported in Fig. 2, that is, 10^3^ simulations of *N* = 50 individuals that have individual accuracy *α* drawn from a normal distribution 𝒩 (*μ*_*α*_ = 0.5, *σ*_*α*_) (flipping to 1−*α* when *α <* 0.5), and varying heterogeneity *σ*_*α*_ (indicated next to the curve). Shaded areas are 95% confidence intervals. We show the results for communication on (a) Erdos-Renyi (ER) graphs with link probability *p* = 0.2 and (b) Barabasi-Albert (BA) networks with number of edges *m* = 3. Via synchronous updates of individuals’ confidence through the Weighted Bayes Consensus rule (WBC, blue lines) the group reaches a consensus in less time than the Belief Consensus algorithm (BC, green lines); however with lower accuracy (computed as the proportion of runs with unanimous agreement for *S*^+^). The results for ER graphs show a smaller difference between the two strategies than the results of Fig. 2 for random geometric graph (RGG), possibly due to a lower clustering coefficient of ER graphs than RGG. We can also note that BA networks, characterised by highly connected nodes and a scale-free edge distribution, lead to larger differences between strategies than ER graphs which have an approximately homogeneous edge distribution (*i.e*. every node has on average *p N* = 10 neighbours).

**Figure SF4:**
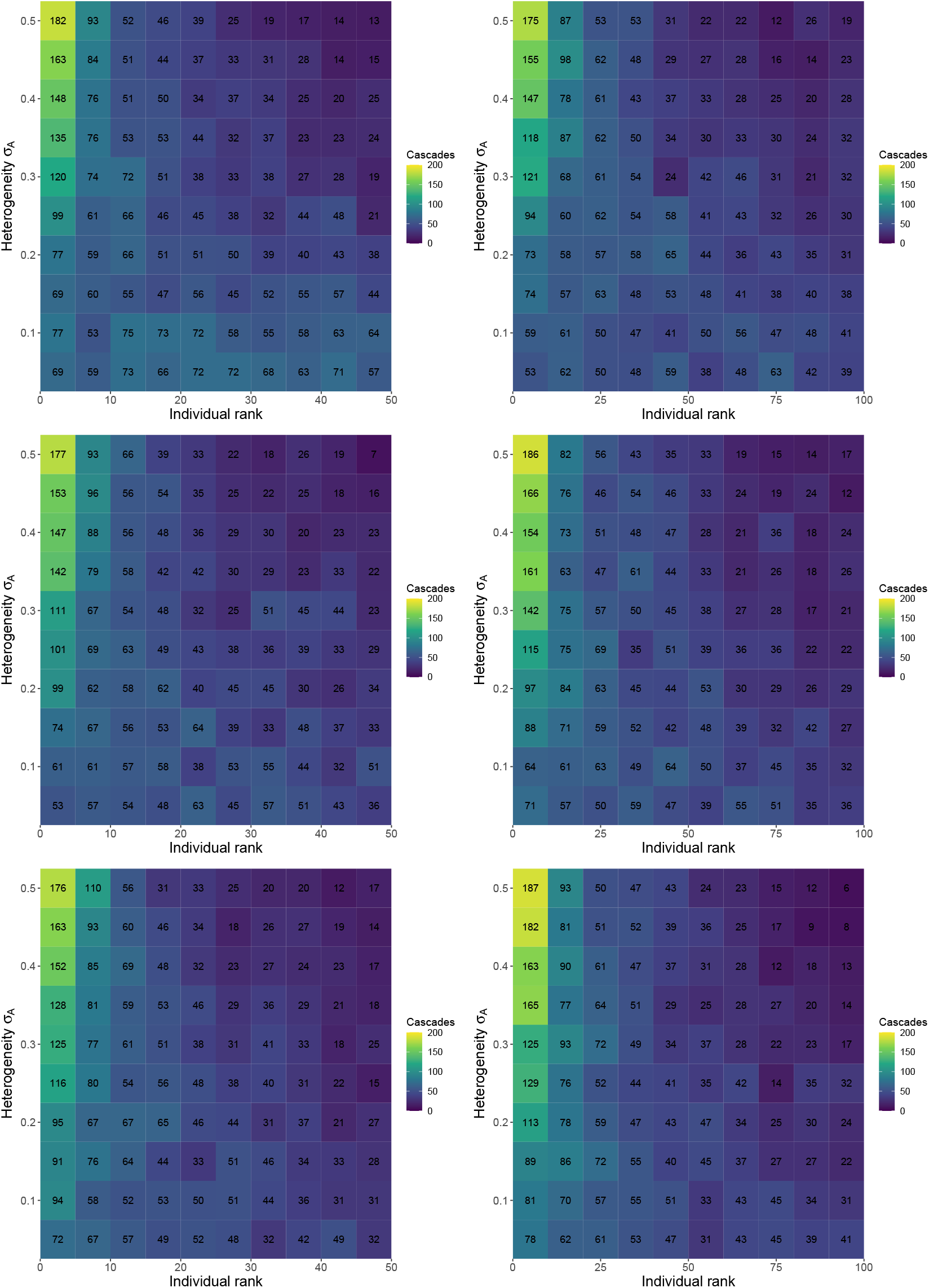
The results of Fig. 3(b) in the main text generalise to Erdos-Renyi (ER) graphs for different group sizes, that is, in heterogeneous groups, the best individuals— with higher drift/noise ratio, *Ã/σ*—more often trigger a cascade. We sort (on the x-axis) the individuals in decreasing order of drift/noise ratio and report the number of cascades each individual triggers. We count as a cascade the triggering of a sequence of at least *N/*10 decisions. The results are from 500 simulation runs of a group of *N* = 50 individuals (left column), of *N* = 100 individuals (right column), communicating on a ER graph with link probability *p* = 0.2, and with drift sampled from 𝒩 (*μ*_*A*_, *σ*_*A*_). The mean drift *μ*_*A*_ is *μ*_*A*_ = 0.1 in the first row, *μ*_*A*_ = 0.2 in the second row, and *μ*_*A*_ = 0.3 in the third row; instead the value of *σ*_*A*_ is indicated on the y-axis. In all tested conditions, the results are qualitatively the same of the ones in Fig. 3(b).

**Figure SF5:**
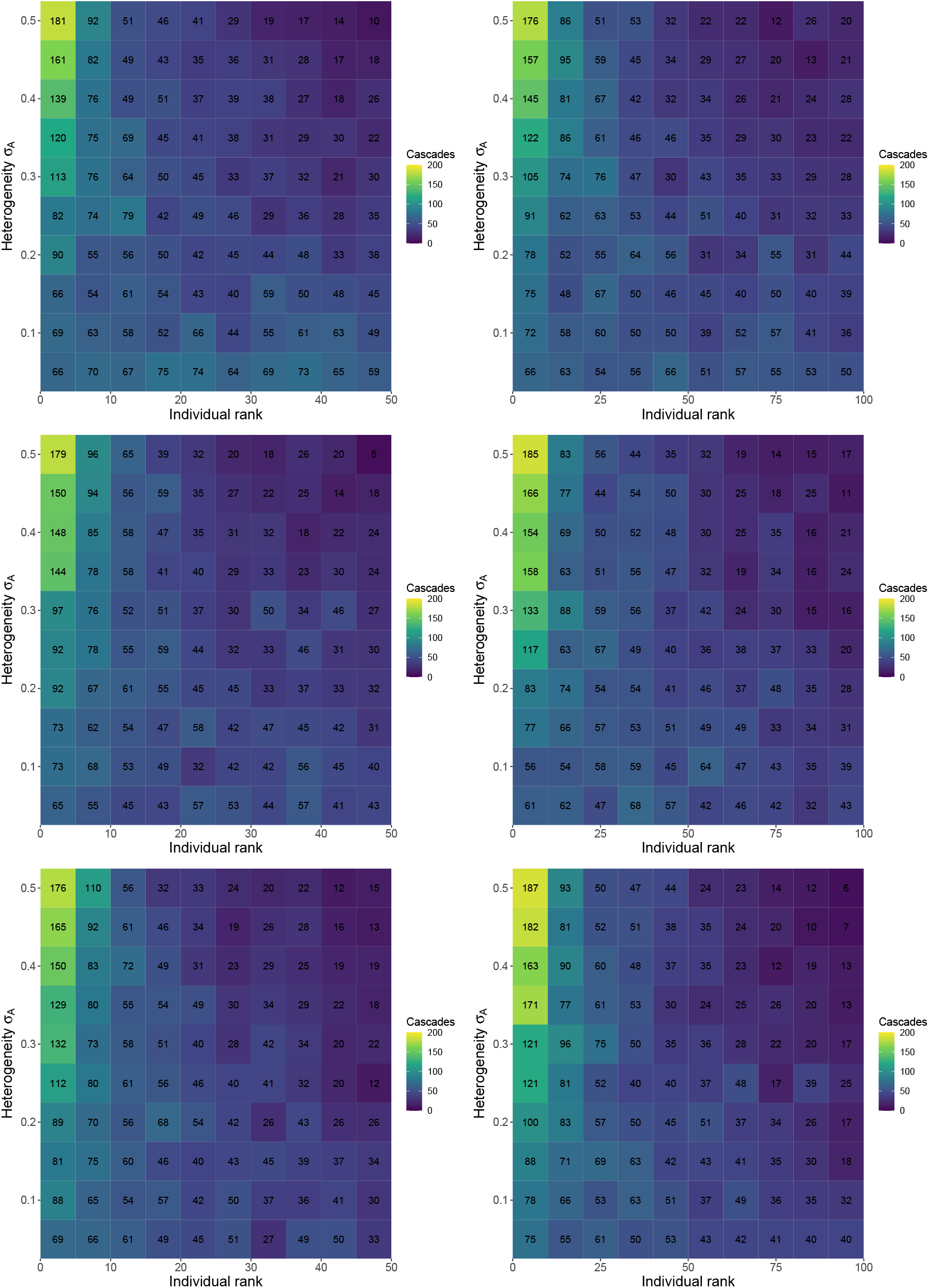
Results for simulation similar to the ones of Fig. SF4, with the difference that here individuals communicated on a Barabasi-Albert (BA) network with number of edges *m* = 5. The results are from 500 simulation runs of a group of *N* = 50 individuals (left column), of *N* = 100 individuals (right column) with drift sampled from 𝒩 (*μ*_*A*_, *σ*_*A*_). The mean drift *μ*_*A*_ is *μ*_*A*_ = 0.1 in the first row, *μ*_*A*_ = 0.2 in the second row, and *μ*_*A*_ = 0.3 in the third row; instead the value of *σ*_*A*_ is indicated on the y-axis. In all tested conditions, the results are qualitatively the same of the ones in Fig. 3(b).

**Figure SF6:**
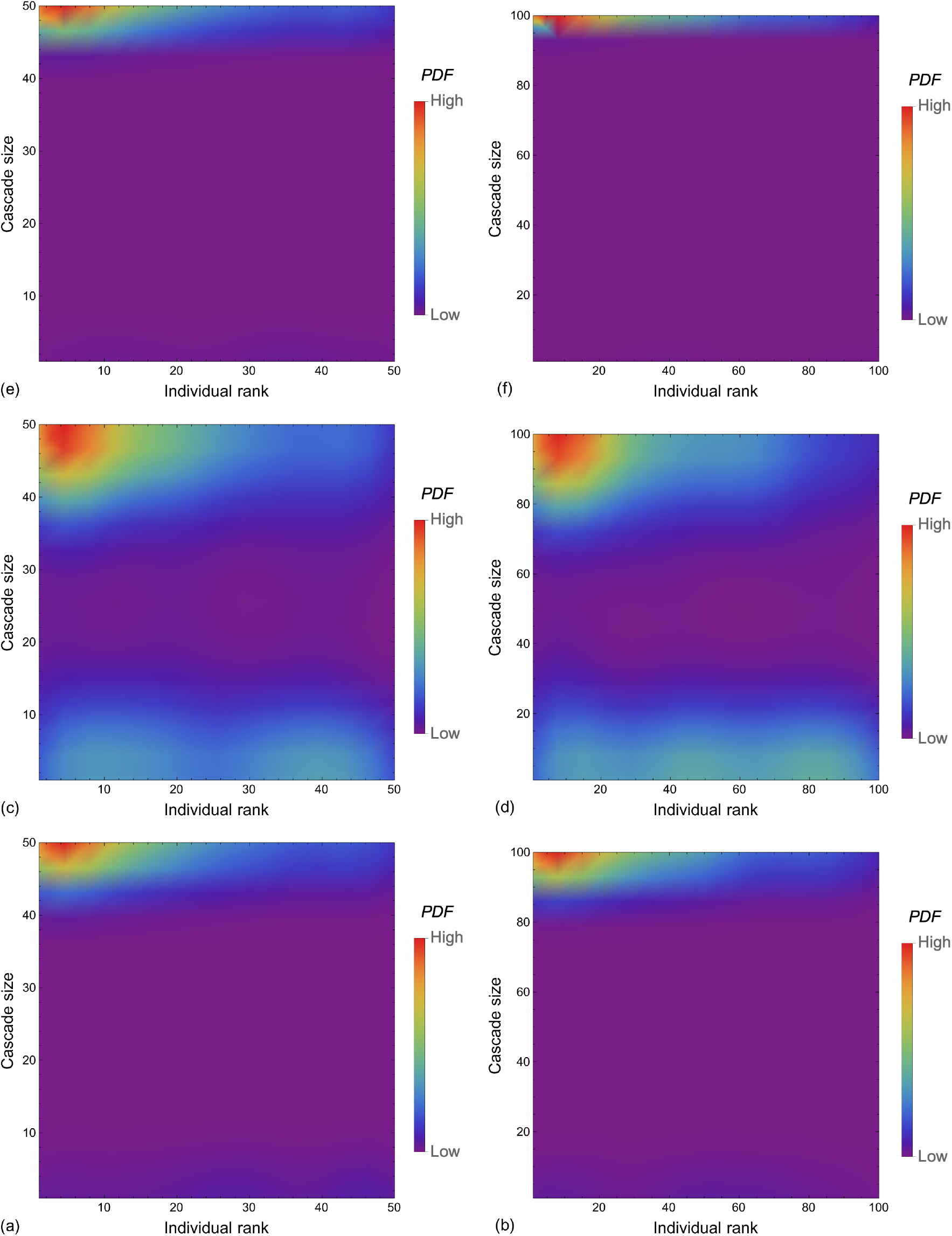
The most accurate individuals trigger the largest cascades in different network topologies. We show the probability density function for each individual, sorted in decreasing drift/noise ratio order (on the x-axis), to trigger a cascade of different sizes (on the y-axis). We count as a cascade the triggering of a sequence of at least *N/*10 decisions. The PDF is computed from 500 simulation runs of a group of *N* = 50 individuals in panels (a,c,e), and *N* = 100 individuals in panels (b,d,f), with drift sampled from 𝒩 (*μ*_*A*_ = 0.2, *σ*_*A*_ = 0.05). We show the results for individuals communicating on a sparse network of three different types: (a-b) Erdos-Renyi with link probability *p* = 0.2, (c-d) random geometric graph with average degree *κ* = 10, and (e-f) Barabasi-Albert with number of edges *m* = 5.

## Notes

https://github.com/DiODeProject/DecisionsOnNetworks

https://doi.org/10.5281/zenodo.7032373

### Competing Interest Statement

The authors have declared no competing interest.

### Summary of Updates

Improved presentation and discussion.

## References

[1] S. Bikhchandani, D. Hirshleifer, and I. Welch, “A theory of fads, fashion, custom, and cultural change as informational cascades,” Journal of Political Economy, vol. 100, no. 5, pp. 992–1026, 1992.

[2] A. V. Banerjee, “A simple model of herd behavior,” The Quarterly Journal of Economics, vol. 107, no. 3, pp. 797–817, 1992.

[3] L. Smith and P. Sorensen, “Pathological outcomes of observational learning,” Econometrica, vol. 68, no. 2, pp. 371–398, 2000.

[4] B. Çelen and S. Kariv, “Observational learning under imperfect information,” Games and Economic Behavior, vol. 47, no. 1, pp. 72–86, 2004.

[5] P. S. Laplace, Theorie Analytique des Probabilites. Paris: Ve Courcier, 1812.

[6] D. C. Knill and A. Pouget, “The Bayesian brain: the role of uncertainty in neural coding and computation,” Trends in Neurosciences, vol. 27, no. 12, pp. 712–719, 2004.

[7] J. M. McNamara, R. F. Green, and O. Olsson, “Bayes’ theorem and its applications in animal behaviour,” Oikos, vol. 112, no. 2, pp. 243–251, 2006.

[8] K. Friston, “The free-energy principle: a unified brain theory?,” Nature Reviews Neuroscience, vol. 11, no. 2, pp. 127–138, 2010.

[9] P. C. Trimmer, A. I. Houston, J. A. Marshall, M. T. Mendl, E. S. Paul, and J. M. McNamara, “Decision-making under uncertainty: biases and bayesians,” Animal Cognition, vol. 14, no. 4, pp. 465–476, 2011.

[10] A. Pouget, J. M. Beck, W. J. Ma, and P. E. Latham, “Probabilistic brains: knowns and unknowns,” Nature Neuroscience, vol. 16, no. 9, p. 1170, 2013.

[11] J. A. Marshall, G. Brown, and A. N. Radford, “Individual confidence-weighting and group decision-making,” Trends in Ecology & Evolution, vol. 32, no. 9, pp. 636–645, 2017.

[12] B. Bahrami, K. Olsen, P. E. Latham, A. Roepstorff, G. Rees, and C. D. Frith, “Optimally interacting minds,” Science, vol. 329, no. 5995, pp. 1081–1085, 2010.

[13] M. Penrose, Random Geometric Graphs. No. 5, Oxford University Press, 2003.

[14] R. Ratcliff, P. L. Smith, S. D. Brown, and G. McKoon, “Diffusion decision model: Current issues and history,” Trends in Cognitive Sciences, vol. 20, no. 4, pp. 260–281, 2016.

[15] R. Kiani and M. N. Shadlen, “Representation of confidence associated with a decision by neurons in the parietal cortex,” Science, vol. 324, no. 5928, pp. 759–64, 2009.

[16] R. Bogacz, E. Brown, J. Moehlis, P. Holmes, and J. D. Cohen, “The physics of optimal decision making: A formal analysis of models of performance in two-alternative forced choice tasks,” Psychological Review, vol. 4, no. 113, pp. 700–765, 2006.

[17] R. Ratcliff, “A theory of memory retrieval,” Psychological review, vol. 85, no. 2, p. 59, 1978.

[18] R. Ratcliff and G. McKoon, “The diffusion decision model: theory and data for two-choice decision tasks,” Neural computation, vol. 20, no. 4, pp. 873–922, 2008.

[19] T. Bayes, “An essay towards solving a problem in the doctrine of chances. by the late rev. mr. bayes, frs communicated by mr. price, in a letter to john canton, amfr s,” Philosophical transactions of the Royal Society of London, vol. 53, pp. 370–418, 1763.

[20] B. Karamched, S. Stolarczyk, Z. P. Kilpatrick, and K. Josić, “Bayesian evidence accumulation on social networks,” SIAM Journal on Applied Dynamical Systems, vol. 19, no. 3, pp. 1884–1919, 2020.

[21] S. Nitzan and J. Paroush, “Optimal decision rules in uncertain dichotomous choice situations,” International Economic Review, vol. 23, no. 2, pp. 289–297, 1982.

[22] P. J. Boland, “Majority systems and the condorcet jury theorem,” Journal of the Royal Statistical Society. Series D (The Statistician), vol. 38, no. 3, pp. 181–189, 1989.

[23] P. C. Trimmer, A. I. Houston, J. A. Marshall, R. Bogacz, E. S. Paul, M. T. Mendl, and J. M. McNamara, “Mammalian choices: combining fast-but-inaccurate and slow-but-accurate decision-making systems,” Proceedings of the Royal Society B: Biological Sciences, vol. 275, no. 1649, pp. 2353–2361, 2008.

[24] R. Olfati-Saber, E. Franco, E. Frazzoli, and J. S. Shamma, “Belief consensus and distributed hypothesis testing in sensor networks,” Networked Embedded Sensing and Control, pp. 169–182, 2006.

[25] A. Wald and J. Wolfowitz, “Optimum character of the sequential probability ratio test,” The Annals of Mathematical Statistics, pp. 326–339, 1948.

[26] W. Edwards, “Optimal strategies for seeking information: Models for statistics, choice reaction times, and human information processing,” Journal of Mathematical Psychology, vol. 2, no. 2, pp. 312–329, 1965.

[27] M. H. DeGroot, “Reaching a consensus,” Journal of the American Statistical Association, vol. 69, no. 345, p. 118, 1974.

[28] V. Bala and S. Goyal, “Learning from neighbours,” Review of Economic Studies, vol. 65, no. 3, pp. 595–621, 1998.

[29] V. Bala and S. Goyal, “Conformism and diversity under social learning,” Economic Theory, vol. 17, no. 1, pp. 101–120, 2001.

[30] P. M. DeMarzo, D. Vayanos, and J. Zwiebel, “Persuasion bias, social influence, and unidimensional opinions,” The Quarterly Journal of Economics, vol. 118, no. 3, pp. 909–968, 2003.

[31] C. P. Chamley, Rational herds: Economic models of social learning. Cambridge University Press, 2004.

[32] A. Banerjee and D. Fudenberg, “Word-of-mouth learning,” Games and Economic Behavior, vol. 46, no. 1, pp. 1–22, 2004.

[33] B. Golub and M. O. Jackson, “Naïve learning in social networks and the wisdom of crowds,” American Economic Journal: Microeconomics, vol. 2, no. 1, pp. 112–149, 2010.

[34] M. O. Jackson, “An overview of social networks and economic applications,” in Handbook of Social Economics, vol. 1, pp. 511–585, Elsevier B.V., 2011.

[35] L. Corazzini, F. Pavesi, B. Petrovich, and L. Stanca, “Influential listeners: An experiment on persuasion bias in social networks,” European Economic Review, vol. 56, no. 6, pp. 1276–1288, 2012.

[36] A. Jadbabaie, P. Molavi, A. Sandroni, and A. Tahbaz-Salehi, “Non-Bayesian social learning,” Games and Economic Behavior, vol. 76, no. 1, pp. 210–225, 2012.

[37] P. Ortoleva and E. Snowberg, “Overconfidence in political behavior,” American Economic Review, vol. 105, no. 2, pp. 504–535, 2015.

[38] T. Gagnon-Bartsch and M. Rabin, “Naive social learning, mislearning, and un-learning,” Mimeo, 2016.

[39] E. Mossel and O. Tamuz, “Opinion exchange dynamics,” Probability Surveys, vol. 14, pp. 155–204, 2017.

[40] P. Molavi, A. Tahbaz-Salehi, and A. Jadbabaie, “A theory of non-Bayesian social learning,” Econometrica, vol. 86, no. 2, pp. 445–490, 2018.

[41] G. Levy and R. Razin, “Information diffusion in networks with the Bayesian peer influence heuristic,” Games and Economic Behavior, vol. 109, no. 681579, pp. 262–270, 2018.

[42] D. Acemoglu, M. A. Dahleh, I. Lobel, and A. Ozdaglar, “Bayesian learning in social networks,” The Review of Economic Studies, vol. 78, no. 4, pp. 1201–1236, 2011.

[43] D. Acemoglu, K. Bimpikis, and A. Ozdaglar, “Dynamics of information exchange in endogenous social networks,” Theoretical Economics, vol. 9, no. 1, pp. 41–97, 2014.

[44] E. Mossel, N. Olsman, and O. Tamuz, “Efficient Bayesian learning in social networks with Gaussian estimators,” in 2016 54th Annual Allerton Conference on Communication, Control, and Computing (Allerton), pp. 425–432, IEEE, 2016.

[45] D. Gale and S. Kariv, “Bayesian learning in social networks,” Games and Economic Behavior, vol. 45, no. 2, pp. 329–346, 2003.

[46] E. Mossel and O. Tamuz, “Iterative maximum likelihood on networks,” Advances in Applied Mathematics, vol. 45, no. 1, pp. 36–49, 2010.

[47] G. Levy and R. Razin, “Correlation neglect, voting behavior, and information aggregation,” American Economic Review, vol. 105, no. 4, pp. 1634–1645, 2015.

[48] G. Levy and R. Razin, “Does polarisation of opinions lead to polarisation of platforms? the case of correlation neglect,” Quarterly Journal of Political Science, vol. 10, no. 3, pp. 321–355, 2015.

[49] S. Arganda, A. Pérez-Escudero, and G. G. de Polavieja, “A common rule for decision making in animal collectives across species,” Proceedings of the National Academy of Sciences, vol. 109, no. 50, pp. 20508–20513, 2012.

[50] R. P. Mann, “Collective decision making by rational individuals,” Proceedings of the National Academy of Sciences, vol. 115, no. 44, pp. E10387–E10396, 2018.

[51] L. Wang and F. Xiao, “Finite-time consensus problems for networks of dynamic agents,” IEEE Transactions on Automatic Control, vol. 55, no. 4, pp. 950–955, 2010.

[52] P. Jia, A. MirTabatabaei, N. E. Friedkin, and F. Bullo, “Opinion dynamics and the evolution of social power in influence networks,” SIAM Review, vol. 57, no. 3, pp. 367–397, 2015.

[53] V. Amelkin, F. Bullo, and A. K. Singh, “Polar opinion dynamics in social networks,” IEEE Transactions on Automatic Control, vol. 62, no. 11, pp. 5650–5665, 2017.

[54] M. Ye, Opinion Dynamics and the Evolution of Social Power in Social Networks. Springer International Publishing, 2019.

[55] J. Liu, M. Ye, B. D. Anderson, T. Basar, and A. Nedic, “Discrete-Time Polar Opinion Dynamics with Heterogeneous Individuals,” in 2018 IEEE Conference on Decision and Control (CDC), no. Cdc, pp. 1694–1699, IEEE, 2018.

[56] S. A. Marvel, J. Kleinberg, R. D. Kleinberg, and S. H. Strogatz, “Continuous-time model of structural balance,” Proceedings of the National Academy of Sciences, vol. 108, no. 5, pp. 1771–1776, 2011.

[57] P. Dandekar, A. Goel, and D. T. Lee, “Biased assimilation, homophily, and the dynamics of polarization,” Proceedings of the National Academy of Sciences, vol. 110, no. 15, pp. 5791–5796, 2013.

[58] A. C. R. Martins and S. Galam, “Building up of individual inflexibility in opinion dynamics,” Physical Review E, vol. 87, no. 4, p. 042807, 2013.

[59] C. E. La Rocca, L. A. Braunstein, and F. Vazquez, “The influence of persuasion in opinion formation and polarization,” EPL (Europhysics Letters), vol. 106, no. 4, p. 40004, 2014.

[60] P. Balenzuela, J. P. Pinasco, and V. Semeshenko, “The undecided have the key: Interaction-driven opinion dynamics in a three state model,” PLOS ONE, vol. 10, no. 10, p. e0139572, 2015.

[61] J. P. Pinasco, V. Semeshenko, and P. Balenzuela, “Modeling opinion dynamics: Theoretical analysis and continuous approximation,” Chaos, Solitons & Fractals, vol. 98, pp. 210–215, 2017.

[62] A. Woolcock, C. Connaughton, Y. Merali, and F. Vazquez, “Fitness voter model: Damped oscillations and anomalous consensus,” Physical Review E, vol. 96, no. 3, p. 032313, 2017.

[63] V. Srivastava and N. E. Leonard, “Collective decision-making in ideal networks: The speed-accuracy tradeoff,” IEEE Transactions on Control of Network Systems, vol. 1, no. 1, pp. 121–132, 2014.

[64] B. Karamched, M. Stickler, W. Ott, B. Lindner, Z. P. Kilpatrick, and K. Josić, “Heterogeneity improves speed and accuracy in social networks,” Physical Review Letters, vol. 125, no. 21, p. 218302, 2020.

[65] J. Morand-Ferron and J. L. Quinn, “Larger groups of passerines are more efficient problem solvers in the wild,” Proc. Natl. Acad. Sci. U.S.A., vol. 108, no. 38, p. 15898, 2011.

[66] C. C. Ioannou, “Swarm intelligence in fish? the difficulty in demonstrating distributed and self-organised collective intelligence in (some) animal groups,” Behav. Process., vol. 141, p. 141, 2017.

[67] I. D. Couzin, “Collective cognition in animal groups,” Trends in Cognitive Sciences, vol. 13, no. 1, pp. 36–43, 2009.

[68] L. R. Anderson and C. A. Holt, “Information cascades in the laboratory,” The American Economic Review, vol. 87, no. 5, pp. 847–862, 1997.

[69] L. Giraldeau, T. J. Valone, and J. J. Templeton, “Potential disadvantages of using socially acquired information,” Philosophical Transactions of the Royal Society of London. Series B: Biological Sciences, vol. 357, no. 1427, pp. 1559–1566, 2002.

[70] G. Rieucau and L. A. Giraldeau, “Exploring the costs and benefits of social information use: An appraisal of current experimental evidence,” Philosophical Transactions of the Royal Society B: Biological Sciences, vol. 366, no. 1567, pp. 949–957, 2011.

[71] A. J. W. Ward, J. E. Herbert-Read, D. J. T. Sumpter, and J. Krause, “Fast and accurate decisions through collective vigilance in fish shoals,” Proceedings of the National Academy of Sciences, vol. 108, no. 6, pp. 2312–2315, 2011.

[72] J. W. Jolles, N. J. Boogert, V. H. Sridhar, I. D. Couzin, and A. Manica, “Consistent individual differences drive collective behavior and group functioning of schooling fish,” Current Biology, vol. 27, no. 18, pp. 2862–2868.e7, 2017.

[73] M. Piccione and A. Rubinstein, “On the interpretation of decision problems with imperfect recall,” Games and Economic Behavior, vol. 20, no. 1, pp. 3–24, 1997.

[74] I. Kallir and D. Sonsino, “The neglect of correlation in allocation decisions,” Southern Economic Journal, vol. 75, no. 4, pp. 1045–1066, 2009.

[75] E. Eyster and G. Weizsacker, “Correlation neglect in financial decision-making,” SSRN Electronic Journal, 2010.

[76] E. Eyster, M. Rabin, and G. Weizsacker, “An experiment on social mislearning,” SSRN Electronic Journal, 2015.

[77] B. Enke and F. Zimmermann, “Correlation neglect in belief formation,” The Review of Economic Studies, vol. 86, no. 1, pp. 313–332, 2017.

[78] G. C. Williams, Adaptation and Natural Selection: A Critique of Some Current Evolutionary Thought. Princeton University Press, 1966.

[79] K. N. Laland and K. Williams, “Social transmission of maladaptive information in the guppy,” Behavioral Ecology, vol. 9, no. 5, pp. 493–499, 1998.

[80] P. Pongrácz, Á. Miklósi, E. Kubinyi, J. Topál, and V. Csányi, “Interaction between individual experience and social learning in dogs,” Animal Behaviour, vol. 65, no. 3, pp. 595–603, 2003.

[81] J. J. Nocera, G. J. Forbes, and L.-A. Giraldeau, “Inadvertent social information in breeding site selection of natal dispersing birds,” Proceedings of the Royal Society B: Biological Sciences, vol. 273, no. 1584, pp. 349–355, 2006.

[82] G. Rieucau and L.-A. Giraldeau, “Persuasive companions can be wrong: the use of misleading social information in nutmeg mannikins,” Behavioral Ecology, vol. 20, no. 6, pp. 1217–1222, 2009.

[83] A. Avarguès-Weber, R. Lachlan, and L. Chittka, “Bumblebee social learning can lead to suboptimal foraging choices,” Animal Behaviour, vol. 135, pp. 209–214, 2018.

[84] F.-X. Dechaume-Moncharmont, A. Dornhaus, A. I. Houston, J. M. McNamara, E. J. Collins, and N. R. Franks, “The hidden cost of information in collective foraging,” Proceedings of the Royal Society B: Biological Sciences, vol. 272, no. 1573, pp. 1689–1695, 2005.

[85] C. Grüter and E. Leadbeater, “Insights from insects about adaptive social information use,” Trends in Ecology & Evolution, vol. 29, no. 3, pp. 177–184, 2014.

[86] V. Srivastava, S. F. Feng, J. D. Cohen, N. E. Leonard, and A. Shenhav, “A martingale analysis of first passage times of time-dependent Wiener diffusion models,” Journal of Mathematical Psychology, vol. 77, pp. 94–110, 2017.

[87] F. Bullo, Lectures on Network Systems. Kindle Direct Publishing, 1.3 ed., 2019. With contributions by J. Cortes, F. Dorfler, and S. Martinez.

